# Catch-and-Display Immunoassay for Digital Biomarker Detection

**DOI:** 10.64898/2026.01.27.702166

**Authors:** Yuxuan Liu, Sam Walker, Michael Klaczko, Ahmet Gurcan, Benjamin Singer, Michel Godin, Vincent Tabard-Cossa, Jon Flax, James McGrath

## Abstract

Digital immunoassays provide exceptional analytical sensitivity for detecting low-abundance biomarkers, but their broad adoption is limited by practical barriers. Commercial platforms are prohibitively expensive for routine use by individual laboratories, and laboratory-scale concepts typically describe specialized biosensors and sophisticated workflows. Here, we introduce a nanomembrane-based Catch-and-Display Immunoassay (CAD-IA) as a cost-effective and laboratory-friendly digital immunoassay for common research settings. In CAD-IA, fluorescent nanoparticles are “captured” by the nanoscale pores of ultrathin silicon nitride membranes through a pipette powered filtration. The captured nanoparticles serve as optically isolated ‘hotspots’ for fluorescent immunocomplex formation when target antigen is present. Co-localization of the fluorescent particles and fluorescent immunocomplexes are then “displayed” and quantified by standard confocal microscopy to generate digital signals. CAD-IA is implemented using the µSiM-DX (**micro**fluidic device featuring an ultrathin **si**licon **m**embrane for **diagnostics**) platform, which is manually assembled from mass produced, cost-effective components. Using the traumatic brain injury (TBI) biomarker S100B as a model, we demonstrate that CAD-IA provides consistent digital outputs and linear quantification with a dynamic range of at least two orders of magnitude when digital and analog analysis are combined on the same image sets. We further demonstrate that the assay maintains linearity in serum matrices and achieves suitable sensitivity (LoD = 0.02 μg/mL) for clinically relevant diagnostic with the addition of tyramide signal amplification (TSA). While further optimization of CAD-IA is possible, these results constitute a proof-of-concept demonstration of a novel digital immunoassay deployable in most research laboratories.

## Introduction

Digital diagnostic immunoassays have emerged as next-generation platforms for ultrasensitive disease-related biomarker detection for critical clinical intervention^1-4^. In contrast to conventional analog immunoassays, which measure bulk signal intensity, digital formats compartmentalize individual binding events and output binary signals (positive/negative), thereby enabling molecular-level detection with reduced instrumentation noise and significantly improved sensitivity^5-8^. A prime example is the SiMoA (Single Molecule Array) technology developed by Quanterix, which achieves femtomolar sensitivity to protein biomarkers by confining immunocomplex-labeled magnetic nanoparticles in femtoliter wells for digital signal development and readout^9-11^. The Simoa HD-X platform achieves fg/mL level detection limits, but relies on dedicated proprietary instrumentation with capital costs of several hundred thousand dollars, as well as proprietary reagents and consumables. More broadly, commercialized digital immunoassays are often built on specialized instruments featuring high-end detection system, automated microfluidic handlers, and proprietary signal acquisition modules, making them prohibitively expensive for individual research laboratories and low resource environments^12-16^.

Laboratory-manufactured digital immunoassays have been explored as more cost-effective alternatives for use in resource-limited settings. For example, He et al. developed a solid-state nanopore-based digital immunoassay capable of quantifying biomarkers by generating binary readouts from DNA reporter structural transformations^5^. Ren et al. further advanced this concept by integrating click chemistry with a computer-vision-assisted digital immunosensor that enables smartphone-based imaging of immunocomplexes^17^. However, these laboratory-built platforms still require specialized scientific skills to operate or rely on complex fabrication of customized biosensors. In addition, home-built detection hardware and data analysis pipelines can require extensive calibration and parameter tuning to achieve high-quality signals^18-20^. The lack of user-friendly workflows and plug-and-play operation limits the adoption of these solutions by external laboratories.

The µSiM-DX (**micro**fluidic device featuring an ultrathin **si**licon **m**embrane for **diagnostics**) is a microfluidic platform featuring ultrathin silicon membranes advanced by our laboratory over two decades^21-24^. The pipette-powered device comprises a 10 µL microfluidic input channel and a 100 µL exit well separated by a 100-nm-thick nanoporous silicon nitride (NPN) membrane (10^10^ ∼60 nm pores per cm^2^; **Figure 1A**). All components of the µSiM-DX are commercially available and can be manually assembled within minutes, eliminating the need for complex biosensor fabrication^25^. We have previously reported the use of the µSiM-DX for viral detection^24^ and the Catch-and-Display Liquid Biopsy (CAD-LB) for biomarkers assessment on extracellular vesicles (EV)^26^. In CAD-LB, individual EVs are fluorescently tagged with antibodies against disease-associated surface biomarkers and captured on a nanomembrane. The “captured” EVs can then be “displayed” by confocal microscopy and digitally analyzed to enable diagnostic evaluation. Motivated by the success of CAD-LB, we now extend the ‘catch and display’ modality to soluble protein detection with a Catch-and-Display Immunoassay (CAD-IA). Both CAD-LB and CAD-IA are “digital assays” in that they create spatially isolated countable signals in the presence of target analyte. The NPN nanopores allow small molecules to freely pass through, reducing nonspecific background signal and facilitating the efficient targeting of reagents to the surfaces of immobilized isolated nanoparticles (**Figure 1B**). In contrast to the ex situ labeling workflow used in CAD-LB, CAD-IA immobilizes fluorescent nanoparticles on the membrane as “hotspots” and employs an in situ labeling strategy by constructing sandwich immunocomplexes directly on the nanoparticle surfaces through the sequential addition of capture antibody, biomarker-containing samples, and detection antibody labeled with spectrally distinct fluorochromes. Digital measurements are obtained with confocal imaging of the nanomembrane and quantification of colocalization of immunocomplexes and nanoparticles hotspots at different fluorescence wavelengths (**Figure 1C**). Complementary analog measurements are also acquired by quantifying the immunocomplex fluorescence intensity at each nanoparticle hotspot to extend the effective dynamic range of the assay. This unique CAD-IA workflow distinguishes from other nanoparticle-based digital platforms by combining nanomembrane capture, in situ immunocomplex formation, and direct dual digital-plus-analog readout in one streamlined catch–bind–display process^27, 28^.

**Figure 1.**
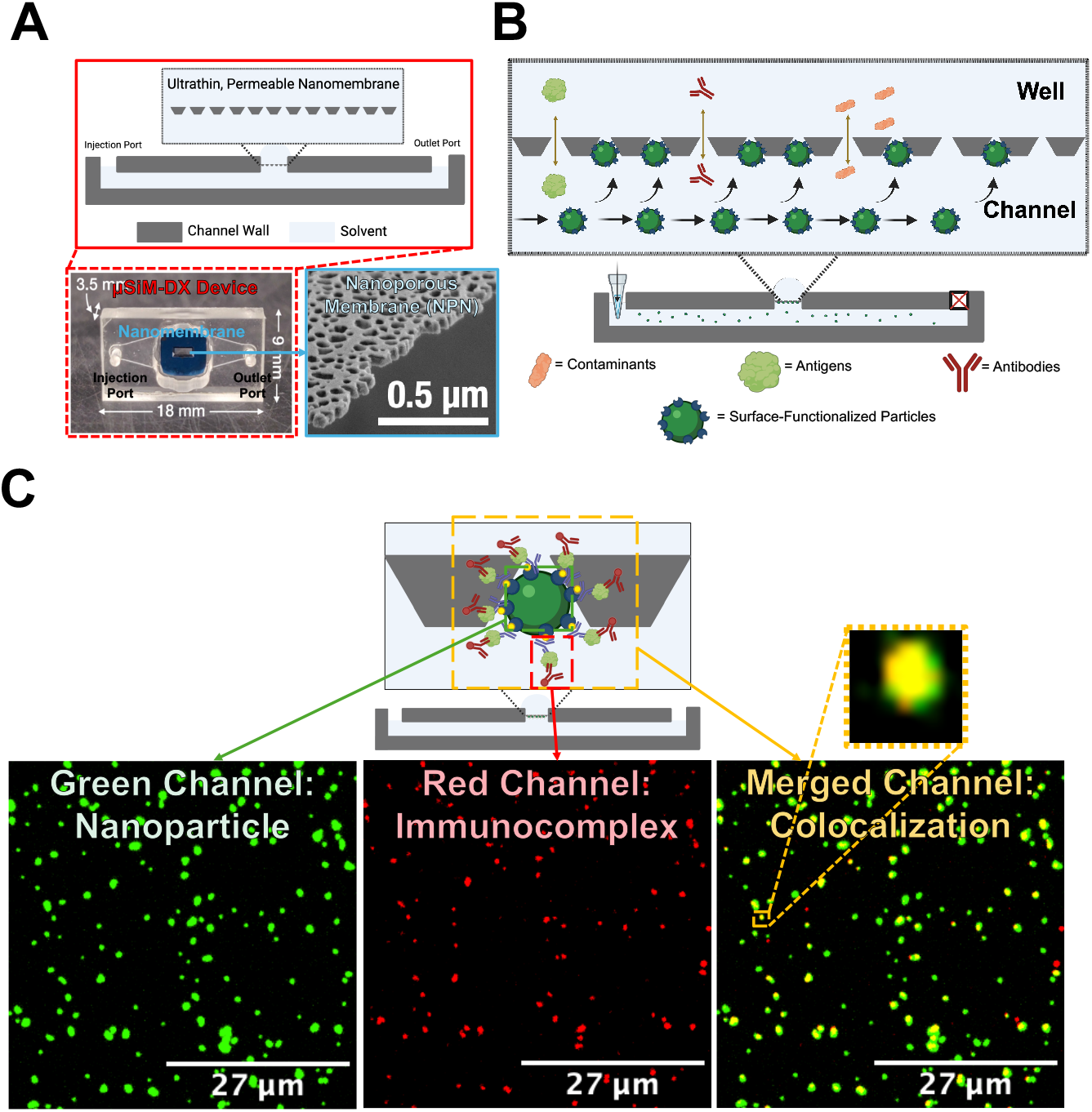
Overview of the Nanomembrane-Based Immunoassay Platform. **(A)** Ultrathin, permeable nanomembrane-integrated μSiM-DX device, consisting of a well and a microfluidic channel with inlet and outlet ports, separated by a nanoporous membrane (red dashed box). The NPN contains nanopores of approximately 60 nm in diameter (blue solid box). **(B)** Conceptual illustration of surface-functionalized nanoparticle capture and molecular transport within the μSiM-DX device. **(C)** Digital signal readout strategy. Positive signals are detected by the colocalization of immunocomplexes (red channel, middle panel) with captured nanoparticles (left panel), as shown in the merged fluorescence image (right panel).

As a generalizable immunoassay platform for soluble protein biomarker detection, we demonstrate the feasibility of CAD-IA in the context of traumatic brain injury (TBI) and use its associated biomarkers as proof-of-principle analytes. TBI is a disruption of normal brain function caused by external mechanical head trauma^29-32^, represents a significant global health burden, affecting approximately 50–60 million individuals annually and incurring an estimated economic cost of ∼ $400 billion worldwide^33^. As TBI can be diagnosed using its associated serum biomarkers, including GFAP and S100B^34-39^, the choice of TBI provides two strategic advantages for assay development. First, these biomarkers typically remain at low baseline levels in healthy individuals, enabling a naturally low background for negative samples and a favorable dynamic window for spiked-in feasibility studies. Second, as well-recognized TBI-associated biomarkers, we are able to employ ELISA-antibody ‘kits’ that are commercially available and well-validated. With these choices, we demonstrate CAD-IA as a novel digital immunoassay for S100B that linearly quantifies S100B from a limit of detection (LoD) of 0.02 µg/mL to a saturation limit of 0.75 µg/mL (1.92 nM to 72 nM, R^2^ = 0.9889). We further show that the same image set used for digital detection can be used in a complementary analog analysis to extend the linear dynamic range to at least 4 µg/mL (384 nM). Finally, we demonstrate that the LoD for S100B can be improved to at least 0.2 ng/mL (0.0192 nM) using tyramide signal amplification (R^2^ = 0.9633)^40-42^, and that CAD-IA is capable of detecting S100B spiked into serum at clinically relevant concentrations. Though CAD-IA involves a tradeoff in sensitivity at the ng/mL level, the platform only costs $35 per assay, which is three to four orders of magnitude less expensive than the Quanterix Simoa HD-X. It is also compatible with high-resolution imaging systems such as confocal microscopes, which are commonly available as shared core facility instruments at many research institutions. Our findings encourage further development of CAD-IA as a cost-effective and laboratory-friendly digital immunoassay and eventual adoption in clinical diagnostics.

## Experimental

### Materials

The following reagents and materials were used in this study: anti-S100B (TA700560 and TA600560), anti-GFAP antibody pairs (TA700041 and TA600039), and S100B (TP300277) were all purchased from Origene. GFAP was purchased from Cayman Chemical Company (27353). Non-PEGylated Dragon Green Fluorescent Polystyrene Nanoparticles (200 nm, Streptavidin-Coated, Ext480/Emi520) were obtained from Bangslaboratory (FSDG002). PEGylated DiagNano™ Green Fluorescent Silica Particles (200 nm, Streptavidin-coated, Ext488/Emi510) were purchased from CD Bioparticles (DNG-L301). The fluorescent detection antibodies were synthesized by conjugating Alexa Fluor 647 using the Alexa Fluor™ 647 Antibody Labeling Kit (A20186, Thermo Fisher Scientific). HRP-conjugated detection antibodies were synthesized using the HRP Conjugation Kit (ab102890, Abcam). The Alexa Fluor 568-labeled anti-mouse secondary antibody was purchased from Thermo (A-11004). Tyramide-labeled Alexa Fluor 647 was purchased from Thermo Fisher Scientific (B40958). 0.1% Bovine Serum Albumin (BSA) and 10% Fetal Bovine Serum (FBS) solution was prepared by dissolving BSA to a final concentration of 0.1% (w/v) and diluting 100% FBS tenfold in phosphate-buffered saline (PBS).

The solution was then ultracentrifuged at 50,000 rpm for 1 hour at 4 °C. After ultracentrifugation, the supernatant (90% of the total volume) was collected and used for subsequent experiments.

### Methods

#### Device Fabrication

µSiM-DX devices were manually assembled in the lab using microfluidic components and device assembly jigs following procedures described in our previous publications^24, 26^. Microfluidic components and assembly jigs were obtained from ALine Inc. (Signal Hill, CA). 1-slot NPN membranes with randomly distributed pores (∼60 nm; ∼ 15% porosity across a 0.014 cm^2^ membrane area) were purchased from SIMPore INC. (SKU: NPSN100-1L). μSiM-DX device assembly has been described in detail previously. Device assembly was performed by applying pressure to acrylic components lined with pressure-sensitive adhesive (PSA) onto membrane chips, with alignment guided by the assembly jigs. Each chip contains a trench side and a flat side surrounding the embedded nanomembrane. The nanomembrane was inserted in the “trench-up” orientation, which exposes more nanoparticle binding sites to the reagents during the assay.

#### Nanoparticle Capture, Reagent Loading, and Washing Procedures within µSiM-DX Device

The capture of nanoparticles was performed following a procedure similar to the one developed for CAD-LB^26^. Briefly, the outlet port of the µSiM-DX was sealed with adhesive tape, and a nanoparticle suspension (∼10^6^ nanoparticles in 40 µL PBS, >60 nm diameter) was introduced through the open inlet using a pipette to allow physical trapping of nanoparticles into the membrane pores. Reagents including antibodies and biomarker-containing samples were loaded by adding 100 µL of reagent solution into the top well, followed by flushing an additional 100 µL through the bottom channel. Unbound reagents and other biocontaminants were removed by washing the well twice with 100 µL of washing buffer and flushing the bottom channel three times with another 100 µL. The excess solution emerging from the outlet during either reagent loading or washing step was absorbed using a Kimtech wipe.

#### Pre-Labeling Workflow

All immunocomplex formation steps were performed in low protein-binding microcentrifuge tubes (88379, Thermo Fisher Scientific). Streptavidin-modified fluorescent nanoparticles were resuspended in 0.1 mL of 2 µg/mL biotinylated capture antibody solution (0.1% BSA in PBS) and incubated for 2 hours at room temperature. The particles were then incubated with 0.1 mL of target TBI biomarker–containing samples for 2 hours. Subsequently, the nanoparticles were incubated with 0.1 mL of 2 µg/mL Alexa Fluor 647-labeled detection antibody solution (0.1% BSA in PBS) for 2 hours to form sandwich immunocomplexes on the nanoparticle surface in the presence of the target biomarker. Finally, the nanoparticles were introduced into the µSiM-DX device for confocal fluorescence imaging. Samples were washed with 1× PBS by centrifugation at 10,000 rpm for 30 minutes for 3 times after each step throughout the procedure.

#### *In situ* Labeling Workflow

Streptavidin-modified fluorescent nanoparticles (in 40 µL PBS, 2.5 × 10^4^ nanoparticles/µL) were first injected and captured in the µSiM-DX device. A 2 µg/mL biotinylated capture antibody solution (0.1% BSA in PBS) was then loaded and incubated for 2 hours, followed by washing to remove unbound antibodies. Target TBI biomarker–containing samples were subsequently introduced into the device and incubated for another 2 hours, followed by an additional washing step. Next, a 2 µg/mL Alexa Fluor 647–labeled detection antibody solution (0.1% BSA in PBS) was added and incubated for 2 hours to complete the formation of fluorescent immunocomplexes in the presence of the bound biomarker. After the final washing step, the µSiM-DX device was subjected to confocal fluorescence imaging. Reagent purification was initially performed using ultracentrifugation to remove large aggregates during process flow development with GFAP. However, this step was later removed due a realization that it wasn’t necessary for S100B detection and a concern that ultracentrifugation would cause losses and reduce signal.

#### Tyramide Signal Amplification

The TSA-mediated CAD-IA followed the same initial three steps (nanoparticle capture, capture antibody immobilization, and biomarker targeting) as the CAD-IA. Fluorescent labeling was performed by first introducing a 2 µg/mL horseradish peroxidase (HRP)-conjugated detection antibody into the µSiM-DX device, followed by a 2-hour incubation. Unbound detection antibodies were then removed by washing with 1× PBS. Subsequently, tyramide-functionalized Alexa Fluor 647 fluorophores (1×) in PBS containing 0.000015% H_2_O_2_ were introduced into the device and incubated for 10 minutes for tyramide signal amplification staining. After staining, unbound fluorophores were thoroughly removed by washing with TRIS NaCl Tween-20 (TNT) buffer (0.1 M Tris-HCl, 0.15 M NaCl, 0.05% Tween-20), followed by additional washing with PBS prior to imaging. As a technical note, we found it helped to optimize the solvents for TSA-enhanced CAD-IA. Specifically, we found that dissolving the capture antibody in pure PBS, while dissolving the HRP-conjugated detection antibody in 0.1% BSA in PBS, preserved both a high reagent immobilization efficiency and excellent assay specificity (**Figure S8**). We also found improved performance if the reagents used for TSA-modified CAD-IA were not ultracentrifuged (**Figure S9**).

#### Fluorescence Imaging

Fluorescence imaging of the nanomembrane and bound analytes was performed within μSiM-DX devices using a spinning disk confocal microscope (Andor Dragonfly) equipped with Zyla 4.2 sCMOS and Sona 2.0B-11 sCMOS cameras. A 60×/1.2 NA water-immersion objective was used to acquire high-resolution images. Excitation was provided by lasers at 405, 488, 561, and 637 nm, in coordination with emission filters of 450/50 nm, 525/50 nm, 600/50 nm, and 700/75 nm, respectively. For imaging, nanoparticles were captured with a 100 ms exposure and 40% laser power. Immunostaining using in situ fluorescence detection antibodies was imaged with 500 ms exposure at 40% power, whereas TSA immunostaining imaging required 1000 ms exposure and 80% laser power. These settings enabled visualization of the fluorescent nanoparticles and immunocomplexes. For statistical analysis, fluorescence images were acquired from 3 to 6 randomly selected fields of view (238 μm × 238 μm) within each μSiM-DX device.

#### Data Analysis

Confocal microscopy images were imported into FIJI (ImageJ) for digital signal analysis and assay sensitivity characterization. Nanoparticles and immunocomplexes were detected using the ComDet v.0.5.5 plugin across relevant fluorescence channels. Detection parameters were defined as follows: approximate particle size, 3 pixels; intensity threshold (in SD), 10 for nanoparticles and 5 for immunocomplexes; and a maximum distance of 6 pixels for identifying colocalized spots. Large particles and immunocomplex aggregates were included and segmented according to the defined pixel size for analysis. Digital readouts of assay sensitivity were quantified as the colocalization percentage, defined as the percentage of nanoparticles colocalized with immunocomplexes relative to the total number of nanoparticles. Digital signals were also evaluated using false positive criteria established in prior CAD-LB studies^26^. The false-positive threshold was defined by the antibody-to-nanoparticle hotspot ratio, which increases linearly^26^. In addition to digital analysis, analog signal quantification was performed by measuring the integrated fluorescence intensity of nanoparticles in the red fluorescence channel, irrespective of their colocalization status. Statistical analysis of the quantitative data was performed using GraphPad Prism. The limit of detection (LoD) was defined as the lowest tested concentration that exhibited a statistically significant difference from the blank.

## Results and Discussions

### Development of the *In situ* Workflow for CAD-IA

The initial design of the CAD-IA followed a similar rationale to traditional bead-based digital ELISA, in which the sample was pre-formed in a test-tube by sequentially adding reagents and biomarkers to create a bead-based immunocomplex prior to capture on NPN in the µSiM-DX (**Figure S1A, see details in Pre-Labeling Workflow from Methods**). The well-recognized TBI biomarker, GFAP^43, 44^ was used as the target in these studies. Unfortunately, results showed significant nanoparticles aggregation occurred during the immunocomplex formation in solution, resulting in poor assay specificity (**Figure S1B and S1C**). We reasoned that this aggregation could arise from multiple sources including: 1) hydrophobic interactions between nanoparticles following surface modifications^45-48^ and 2) the cross-linking of nanoparticles by GFAP which is known to form multimeric structures^49-51^.

To overcome the aggregation seen with the solution-based preparations of the immunocomplex, we explored the possibility of developing an in device (i.e. *in situ*) workflow where the immunocomplex is formed on immobilized nanoparticles pre-captured to the pores of NPN (**Figure 2**). As the first step, we injected 40 µL of 200 nm streptavidin-labeled fluorescent green nanoparticles (2.5 × 10^4^ nanoparticles/µL) into the µSiM-DX and used time-lapse fluorescence microscopy to confirm that nanoparticles monodisperse capture events did occur (**Supplemental Movie M1**). We also used time-lapse microscopy to confirm that pore-captured nanoparticles remained in place despite aggressive washing steps in the bottom channel of the device (**Supplemental Movie M2**). These preliminary studies assured that the streptavidin-conjugated particles: 1) were not pre-aggregated, 2) could be captured individually and 3) could be targeted by subsequent processing steps to build the immunocomplex *in situ*. Note that the capture of streptavidin functionalized beads transforms the nanoporous membrane into a nanostructured surface with physically isolated and optically resolvable target-affinity ‘hotspots’ (**Figure S2**).

**Figure 2.**
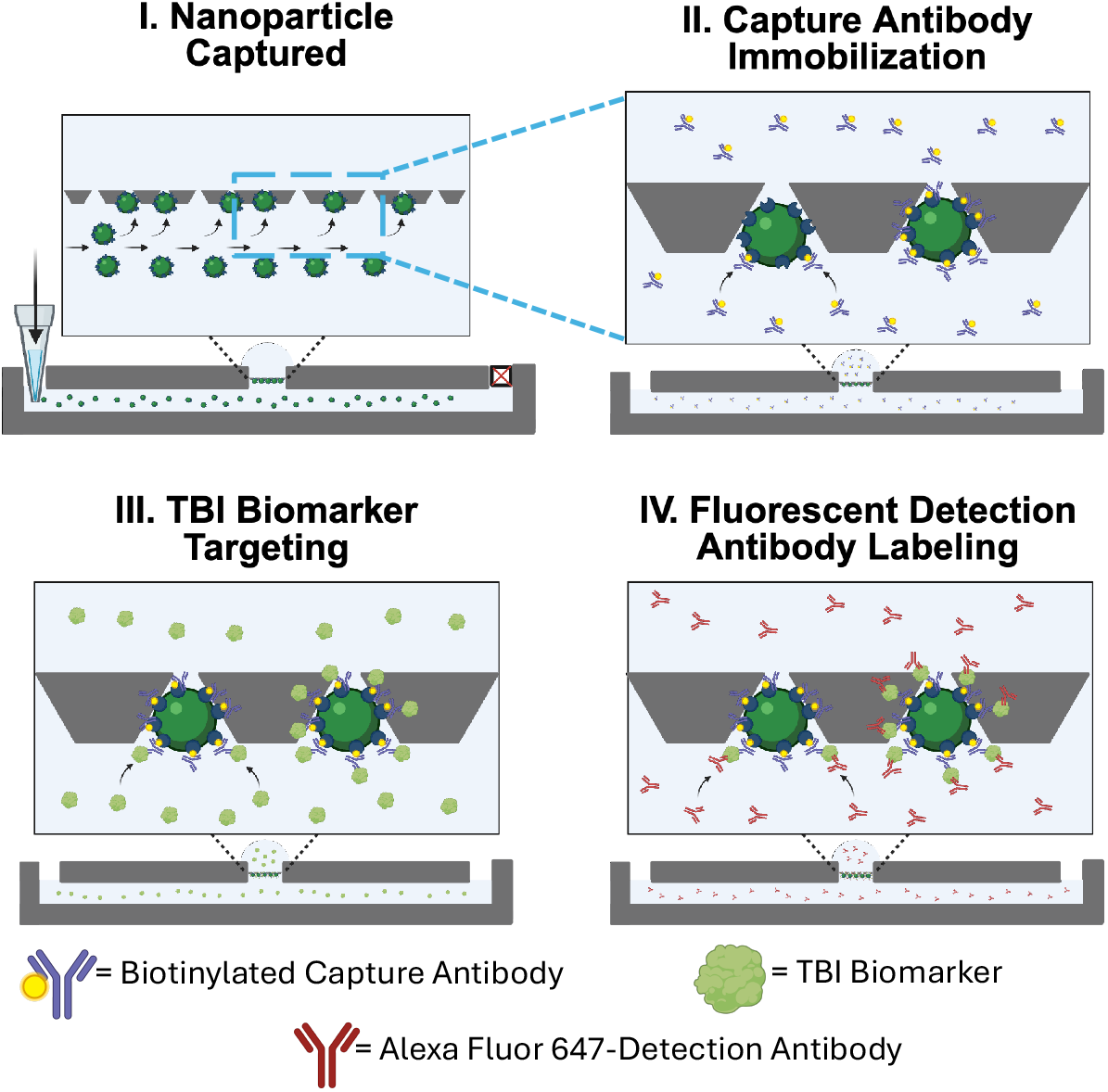
Workflow of the Nanomembrane-Based Catch-and-Display Immunoassay (CAD-IA). **(I)** Streptavidin-labeled nanoparticles captured on the nanomembrane surface. **(II)** Biotinylated capture antibodies immobilized onto the nanoparticle surface via streptavidin–biotin interaction. **(III)** TBI biomarkers bind specifically to the pre-conjugated capture antibodies. **(IV)** Alexa Fluor 647-labeled detection antibodies introduced to form fluorescent immunocomplexes for signal detection.

We next sought to confirm the lack of co-localization with the addition of a fluorescent, non-biotinylated antibody. Without antigen or biotin present in the system, this antibody will have no specificity to the immobilized streptavidin coated nanoparticles and so any co-localization would represent a Type I error (false positive). Surprisingly, nearly 40% of the nanoparticle loci recorded co-localization with the fluorescent antibody. To combat this non-specific binding, we switched to PEGylated streptavidin conjugated nanoparticles, which reduced this nonspecific binding by approximately 400-fold, resulting in only 0.1% colocalization. With this choice we observed very low non-specific binding of the detection antibody (**Figure S4B**).

To establish the feasibility of positive detection events, we performed a second control study where we introduced a non-fluorescent biotinylated primary antibody followed by a secondary fluorescently-tagged antibody with affinity to the primary antibody. Analysis of these experiments demonstrated 92% co-localization of secondary antibody and the nanoparticles (**Figure S5B**). Collectively, these preliminary experiments provide strong evidence of the feasibility of CAD-IA for specific digital detection of binding events on pre-immobilized nanoparticles.

### CAD-IA for S100B Detections

To avoid any potential complications from higher order GFAP structures, we pivoted to a smaller, non-filament forming TBI biomarker S100B^38, 49-54^. We next tested the full CAD-IA *in situ* workflow using S100B as the protein analyte. In these experiments, PEGylated streptavidin-labeled nanoparticles were first captured by NPN, followed by sequential addition and incubation with biotinylated anti-S100B capture antibody, S100B antigen, and Alexa Fluor 647-labeled anti-S100B detection antibody to assemble surface immunocomplexes (**Figure 2**), after which the membrane was imaged by confocal microscopy to generate digital signals of target presence or absence (**see details in *In situ* Labeling Workflow from Methods**). An initial study using S100B dissolved in 0.1% BSA showed consistent co-localization in the presence of analyte (0.2 µg/mL S100B input) and low background in its absence (**Figure 3A**). A dose response study then showed co-localization signal increased with S100B concentrations over the range of 0.02 μg/mL to 0.75 μg/mL (1.92 nM to 72 nM; **Figure 3B Left Panel**) with 0.02 μg/mL (1.92 nM) being the first detectable concentration and thus the LoD. The assay signal reached saturation at S100B concentrations above 0.75 µg/mL (72 nM). Between 0.02 µg/mL and the saturation limit of 0.75 µg/mL CAD-IA showed a strong linear response (R^2^ = 0.9889) (**Figure 3B Right Panel**).

**Figure 3.**
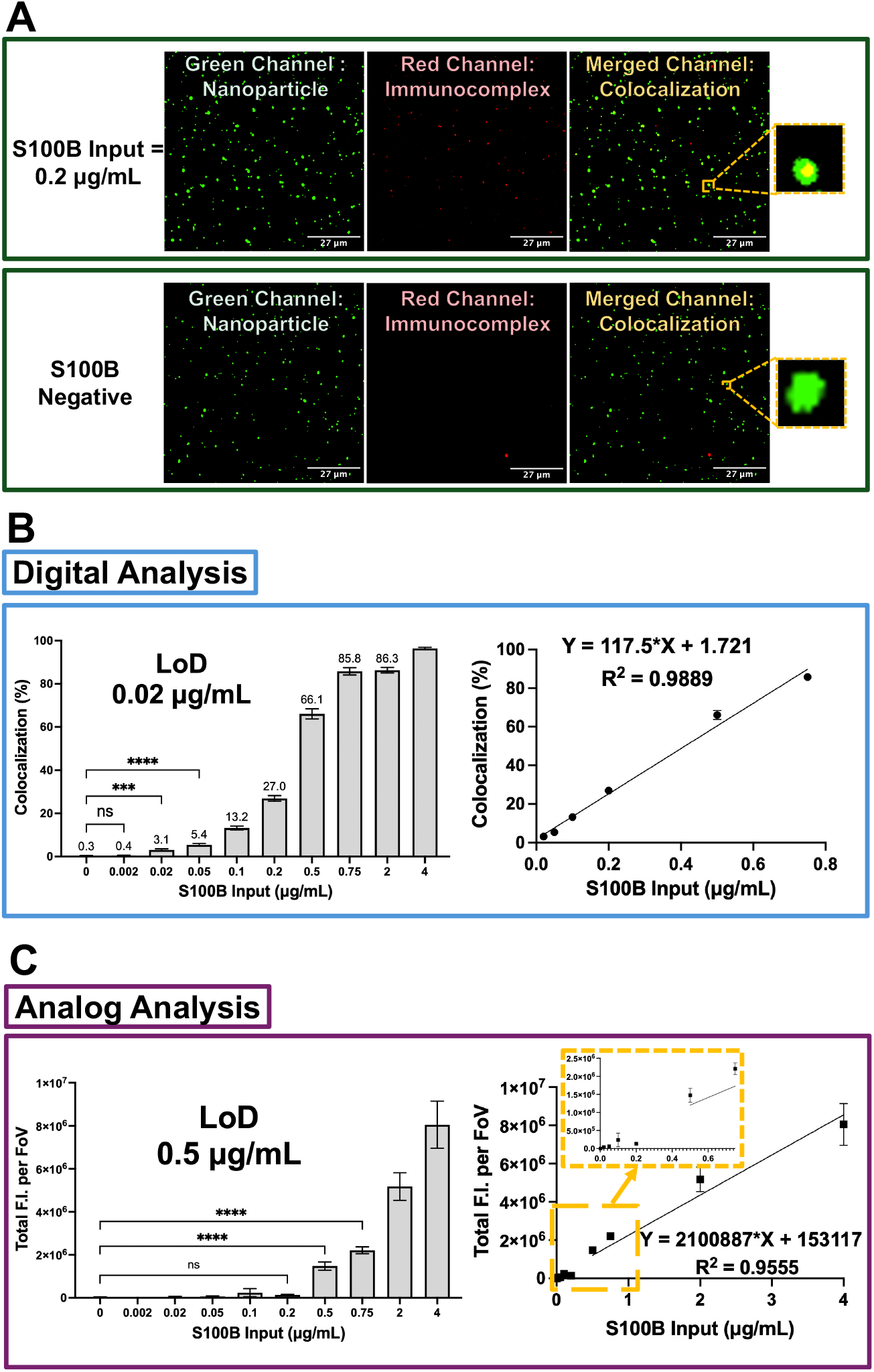
CAD-IA for S100B Detection in 0.1% BSA. **(A)** Representative fields of view for the blank sample (0 µg/mL) and S100B positive (0.2 µg/mL) samples. Pronounced colocalization between immunocomplex fluorescence and nanoparticle hotspots is observed only in the presence of S100B. **(B)** Digital analysis shows dose-dependent response with saturation at 0.75 µg/mL and linear detection from 0.02–0.75 µg/mL (R^2^ = 0.9889). **(C)** Analog analysis based on total fluorescence enables linear quantification beyond 0.5 µg/mL, complementing digital readout and expanding the dynamic detection range. Data are presented as mean ± SD (N = 6 fields of view per device). Statistical significance was evaluated using one-way ANOVA with Dunnett’s test (****p < 0.0001, ***p = 0.0007). Statistical significance is primarily reported for concentrations near the LoD, as higher concentrations exhibited clear separation from the background.

Noticeably, immunocomplex without colocalization to nanoparticle hotspots in merged channel were detected. These signals likely originate from aggregated detection antibodies present in the stock solution or from immunocomplex aggregates formed during the reaction workflow due to incomplete purification. These signals raise the possibility of chance spatial overlap between detection antibody and captured nanoparticles leading to false-positive colocalization. This concern was addressed in the development of CAD-LB^26^ with guidelines for evaluating the frequency of false-positives. Using this method, we estimate a false positive rate by inadvertent co-localization is less than 0.03% indicating that the colocalization we report are true nanoparticle-associated immunocomplex formation.

The sensitivity of digital immunoassays derives from the conversion of target detection events into discrete, countable signals. This is in contrast to analog immunoassays, such as ELISA, which rely on measuring the total signal from bulk binding events which limits sensitivity by increasing signal-to-noise^6, 55, 56^. To confirm these advantages in the context of CAD-IA, we simulated bulk analyses of the same images used for digital analysis in this study by summing the fluorescence intensity of all nanoparticle hotspots in the immunocomplex channel, regardless of their co-localization status. This ELISA-mimicking analysis displays a much higher signal and does not reliably quantify S100B concentrations below 0.5 μg/mL, a fourfold higher LoD than the digital analysis (**Figure 3C Left Panel**).

Interestingly, the analog analysis produces a linear signal response at and above its LoD that extends the detection limit of the assay (**Figure 3C Right Panel**). This suggests that the apparent saturation at 0.75 µg/mL of the digital analysis does not reflect the saturation of S100B binding sites on nanoparticles, but rather is an intrinsic limitation of binary digital analysis, which cannot encode additional increases in signal once all particles are classified as “positive.” The analog and digital analysis of the same images are complementary^6^, with digital assays providing higher sensitivity and analog assays providing extended dynamic range to at least two orders of magnitude. Specifically, low-abundance S100B (0.02 – 0.75 µg/mL) is quantified using the digital mode, wherein discrete nanoparticle-associated immunocomplexes are counted. At higher S100B concentrations (> 0.75 µg/mL), where digital signals become saturated, the assay transitions to an analog mode based on total fluorescence intensity. This dual-readout strategy enables accurate quantification across a wider detectable concentration range, thereby broadening the potential utility of the platform for diverse biomarkers.

Returning to digital CAD-IA, we next tested the ability of the assay to detect S100B in the present of 10% serum in a spike-in assay. We again found the assay produced a consistent and linearly increasing signal with increasing analyte concentration (**Figure S6**). However, the LoD increased from 0.02 μg/mL to 1 μg/mL. This reduced sensitivity can be attributed to the higher background protein concentration in 10% FBS system ( ∼1000 μg/mL in 0.1% BSA to approximately 3000 – 4500 μg/mL in 10% FBS). Despite the presence of PEG on the particle surface, the abundance of albumins and globulins are apparently sufficient to block target-binding sites and hinder the specific interactions required for effective signal generation^57-60^. The loss of sensitivity became complete when S100B was spiked into 100% FBS, where the assay signals dropped to baseline levels across all tested concentrations (**Figure S7**). Hence, for a clinical application of S100B using CAD-IA in the current form, patient serum would need to be diluted at least 10-fold for detection to occur. Normal serum S100B concentrations are approximately 1 × 10^−4^ µg/mL and increases to around 1 × 10^−3^ µg/mL in moderate cases of TBI^61-63^. Even in severe cases, serum-levels of S100B stay at 0.01–0.02 µg/mL in the acute phases of injury^64-66^. Since the projected limit of detection (LoD) of CAD-IA for S100B in serum was estimated to be 10 μg/mL, extrapolated from 1 μg/mL in 10% serum, this sensitivity was insufficient for clinically relevant TBI detection. This result motivated us to explore strategies for further enhancing the sensitivity of CAD-IA.

### Tyramide Signal Amplification (TSA)-Modified CAD-IA for S100B Detections

The need for clinically relevant sensitivity prompted us to explore alternative immunostaining strategies to further improve CAD-IA sensitivity. We designed an modified CAD-IA workflow which incorporate tyramide signal amplification (TSA) into the immunostaining step. TSA is a well-established enzymatic amplification strategy widely used in immunoassay development to enhance detection signals by increasing localized reporter deposition and thereby improve assay sensitivity. The concept of catalyzed reporter deposition (CARD) was first introduced by Robert Bobrow et al., in which an analyte-dependent enzyme, such as horseradish peroxidase (HRP), catalyzes the deposition of labeled phenolic compounds onto nearby solid surfaces, resulting in substantial signal amplification^67^. Subsequent work expanded this concept by exploiting the high reactivity of phenol radicals toward tyrosine residues, which are abundant in proteins, enabling covalent and highly localized reporter accumulation^68^. This mechanism, now commonly referred to as TSA, has been broadly applied to amplify immunoassay signals using HRP-mediated deposition of labeled tyramide molecules^69, 70^. In recent years, TSA has been increasingly adopted to enable the detection of low-abundance analytes that are otherwise challenging to quantify using conventional immunoassays. For example, Choi et al. integrated TSA into a multiplex immunoassay based on encoded hydrogel microparticles, achieving fg/mL level detection of the inflammatory cytokine IL-6, far surpassing the sensitivity of standard ELISA^71^. Similarly, Liu et al. employed a TSA-enhanced ELISA using biotin-tyramide deposition to enable detection of the SARS-CoV-2 spike protein at femtomolar^72^. Akama et al. also introduced a droplet-free bead-based digital ELISA using HRP-catalyzed tyramide deposition confined to individual beads, which inspired our use of TSA to achieve higher sensitivity in CAD-IA^73^.

The TSA-modified CAD-IA preserved the first three steps in the traditional workflow: nanoparticle capture, capture antibody immobilization, and protein biomarker targeting. In the TSA-modified CAD-IA, the detection antibody is pre-conjugated with an enzyme horse radish peroxidase (HRP), rather directly labeled with a fluorophore. As an additional step, tyramide-conjugated Alexa Fluor 647 is then introduced into the µSiM-DX where it is enzymatically activated by particle-localized HRP and covalently links the fluorescent dye to any nearby tyrosine residues in the immunocomplex (**Figure 4, see details in Tyramide Signal Amplification from Methods**). This localized deposition of fluorophores amplifies the detection signal, generating a stronger signal intensity and enhancing assay sensitivity than a fluorophore labeled detection antibody^74, 75^. The TSA modification enhanced the performance of CAD-IA in multiple ways (**Figures 5 and 6**). In both 0.1% BSA and 10% FBS spiked-in assays, the amplified signal showed a consistent S100B dose dependent response over the full dynamic range tested (0 – 1 µg/mL; 0 – 96 nM). Saturation is seen in the case of 10% FBS (at ∼ 0.5 µg /mL; 48nM), but not with 0.1% BSA. Importantly, TSA amplification improved the sensitivity by at least 100-fold for 0.1% BSA (0.0002 µg/mL; 0.0192 nM) and 500-fold in 10% FBS (0.002 µg/mL; 0.192 nM) compared to the unmodified assays (**Figures 5B Left Panel and 6A**). The enzymatic amplification results in a log-linear (**Figures 5B Right Panel and 6B**), rather than linear dose response (**Figure S10**)^76^. This nonlinear dependence arises from enzyme-driven TSA amplification^77^. At low analyte concentrations, catalytic generation of reactive intermediates produces a rapid signal increase, while at higher concentrations the response becomes less steep due to local substrate depletion, radical quenching, and diffusion limitations at high deposition density. More importantly, the TSA enables the assay to achieve a LoD of 0.002 µg/mL in 10% FBS, predicting the successful detection of 0.02 µg/mL in undiluted FBS, achieving a clinically meaningful sensitivity associated with severe cases of TBI^64-66^.

**Figure 4.**
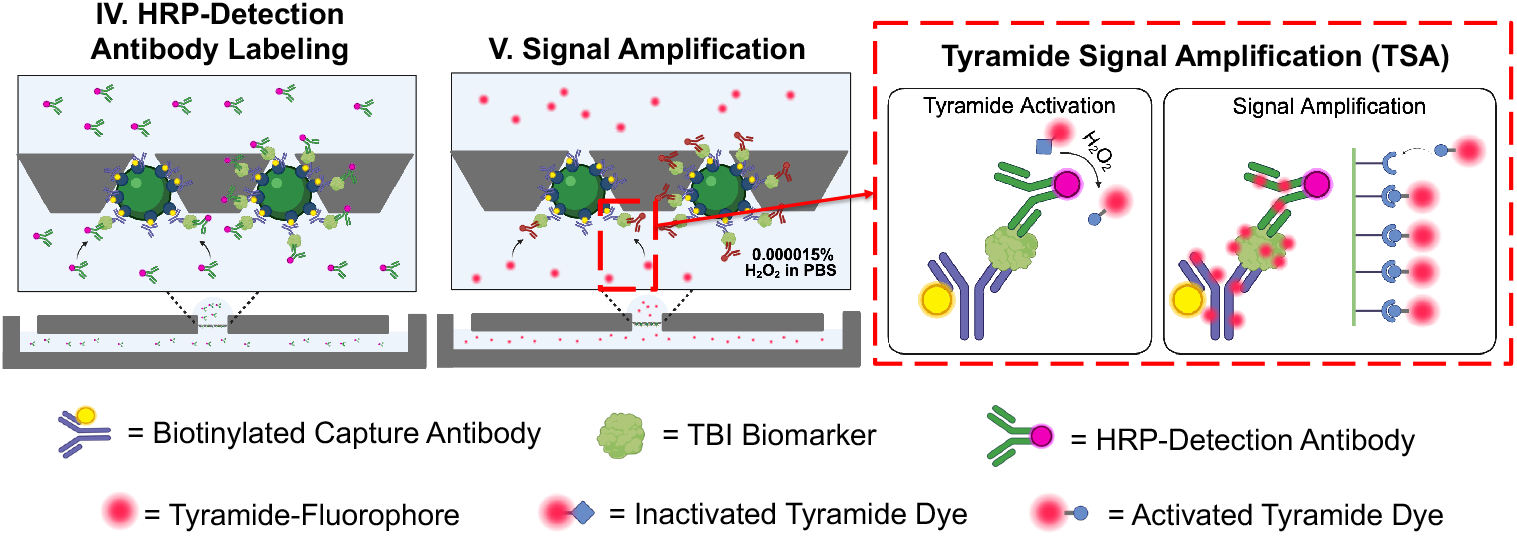
Workflow of TSA-Modified CAD-IA. The TSA-modified CAD-IA preserves the first three steps in the traditional CAD-IA workflow: nanoparticle capture, capture antibody immobilization, and protein biomarker targeting. The fluorescent detection antibody labeling step is replaced by **(IV)** HRP-labeled detection antibodies introduced to form the sandwich complex. **(V)** Tyramide-labeled Alexa Fluor 647 molecules deposited onto tyrosine residues of the immunocomplex through HRP-catalyzed reaction.

**Figure 5.**
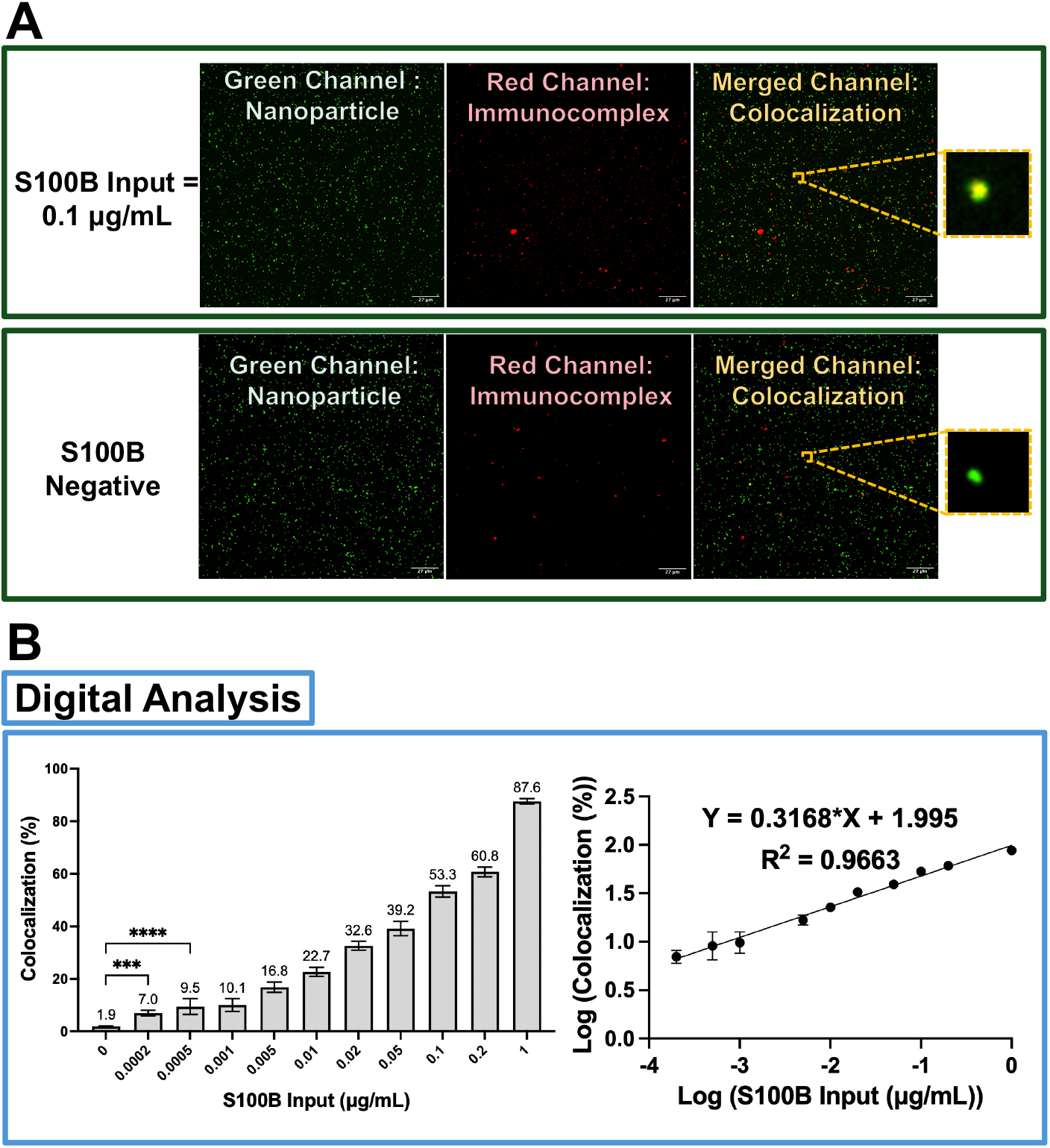
TSA-Modified CAD-IA for S100B Detection in 0.1% BSA. **(A)** Representative fields of view of S100B-negative (no S100B input) and S100B-positive samples (0.1 µg/mL). In the presence of S100B, pronounced colocalization between immunocomplex fluorescence and nanoparticle hotspots is observed, indicating successful formation of target-specific immunocomplexes, whereas negligible colocalization is detected in the absence of S100B. **(B)** Digital Analysis of the TSA-Modified CAD-IA. Dose response analysis demonstrates sensitive and reproducible detection of S100B with a sensitivity down to 0.0002 µg/mL, at least 100-fold higher than CAD-IA not using TSA. Linear regression of the digital signal output shows a strong correlation between colocalization percentage and S100B concentration across the dynamic range of 0.0002–1 µg/mL (R^2^ = 0.9663). Data are presented as mean ± SD (N = 6 fields of view per device). Statistical significance was evaluated using ordinary one-way ANOVA with Dunnett’s multiple comparisons test (****p < 0.0001, ***p = 0.0002).

**Figure 6.**
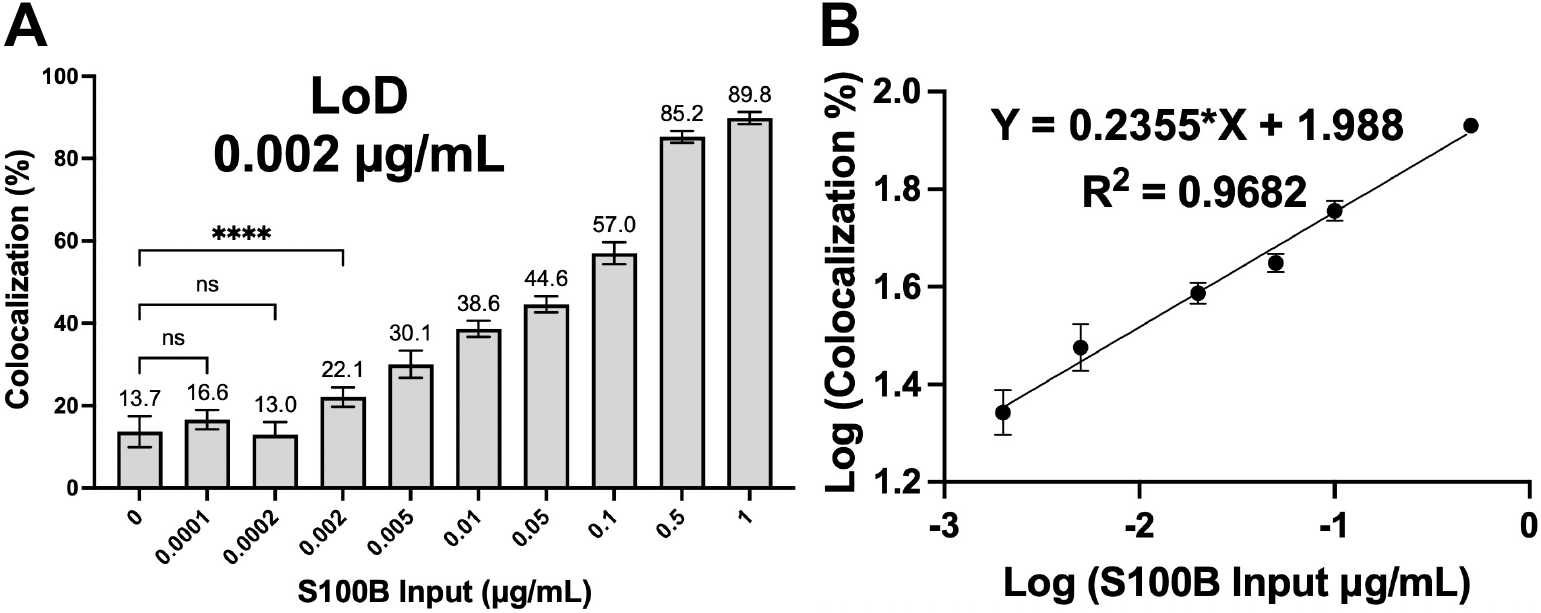
TSA-Modified CAD-IA for S100B Detection in 10% FBS. The assay showed consistent responses with a limit of detection (LoD) of 0.002 μg/mL and a linear range from 0.002 to 1 μg/mL (R^2^ = 0.9682). The projected LoD in 100% FBS is 0.02 μg/mL, exceeding the threshold for severe TBI diagnostic. Data are presented as mean ± SD (N = 6 fields of view per device). Statistical significance was assessed using ordinary one-way ANOVA followed by Dunnett’s multiple comparisons test (****p < 0.0001). Statistical significance is primarily reported for concentrations near the LoD, as higher concentrations exhibited clear separation from the background.

## Conclusions

Here we introduce CAD-IA as a generalizable immunoassay platform for protein detection with cost-effective and commercially accessible microfluidic device (the µSiM-DX). The µSiM-DX features simplified device assembly, and pipette-powered workflow that is adaptable to common research laboratories. CAD-IA uses a specialized *in situ* workflow, in which nanoparticles are firstly immobilized and then functionalized in the pores of an ultrathin nanoporous membrane. The workflow eliminates aggregation artifacts, enables control over the stepwise immunocomplex assembly, and achieves rapid digital readouts with the streamlined art of catch–bind–display. In CAD-IA, immunocomplexes form at physically isolated, optically resolvable, nanopore locations that serve as digital ‘hotspots’ for antigen detection. Using S100B, a representative traumatic brain injury biomarker, as proof-of-principle analyte, we demonstrate its digital detection based on fluorescence colocalization between antigen and nanoparticle, achieving linear quantification at low analyte concentrations with expanded dynamic detection range by the complimentary of analog analysis beyond the detection limit from digital analysis. The assay maintains quantitative performance in serum matrices and incorporates tyramide signal amplification to enable detection at the ng/mL level, encompassing clinically relevant concentrations of S100B. With respect to selectivity, the platform also demonstrates low background and effective suppression of nonspecific binding, as shown by the ∼400-fold reduction in nonspecific colocalization following nanoparticle PEGylation (**Figure S4**).

The value of CAD-IA lies in its ability to convert protein recognition events into spatially resolved, image-based digital signals using minimal sample volume and a simplified workflow. However, further validation and resolution of several remaining limitations are needed prior to broader analytical or clinical deployment. First, systematic evaluation of cross-reactivity against structurally related proteins, including S100A family members and other neurological biomarkers, will be an important focus of future work. This will be paired with studies in spiked serum and CSF matrices to characterize potential matrix effects. Second, sensitivity in complex biological matrices is currently constrained by nanoparticle surface fouling and limitations of the imaging system, which does not yet achieve single-molecule or single-enzyme resolution. Consequently, the observed digital signals arise from concentrated immunocomplex formation rather than true molecular counting. These limitations highlight clear opportunities for improvement through advanced surface chemistries, optimized enzymatic amplification strategies, higher-sensitivity detectors, super-resolution imaging, and automated image-scanning approaches. Third, to expand the broadness of applicable biomarkers in CAD-IA, detection of filament-forming and aggregation-prone antigens such as GFAP should be improved through antigen-specific sample preparation strategies (including mild denaturing protocols) and selection of antibody pairs targeting monomer-specific epitopes. This would enable quantitative definition of the antigen classes for which CAD-IA, in its current form, is best suited, versus those requiring additional optimization. Fourth, the current implementation of CAD-IA has only been demonstrated using confocal microscopy. To improve accessibility, future work should establish workflows compatible with more widely available epifluorescence microscopy platforms. Importantly, the nanopore-localized hotspot architecture is inherently compatible with automation, multiplexing, and high-throughput imaging, enabling simultaneous detection of multiple biomarkers through spectral or spatial encoding. Thus, continued refinement of imaging modalities, assay kinetics, surface engineering, and analytical specificity could enhance the diagnostic utility of CAD-IA for clinically relevant biomarkers and extend the catch-and-display paradigm to a broader range of analytes.

## Author Contributions

Yuxuan Liu contributed to conceptualization, methodology, investigation, data curation, formal analysis, validation, visualization, and writing of the original draft. Samuel Walker and Michael Klaczko contributed to writing – review & editing. Ahmet Gurcan contributed to investigation (SEM imaging) and visualization. Benjamin Singer contributed to conceptualization and writing – review & editing. Michel Godin and Vincent Tabard-Cossa contributed to conceptualization, funding acquisition, and writing – review & editing. Jon Flax and James McGrath contributed to conceptualization, supervision, funding acquisition, and writing – review & editing. All authors reviewed and approved the final manuscript.

## Acknowledgements

The authors gratefully thank the members of the McGrath laboratory and nanomembrane research group for their assistance, feedback, and collaborative environment. This work was supported by NIH RO1EB03158. We further acknowledge the use of shared facilities and instrumentation provided by the Department of Biomedical Engineering at the University of Rochester.

