## Supplementary material for "Catch-and-Display Immunoassay for Digital Biomarker Detection": Supplements.pdf

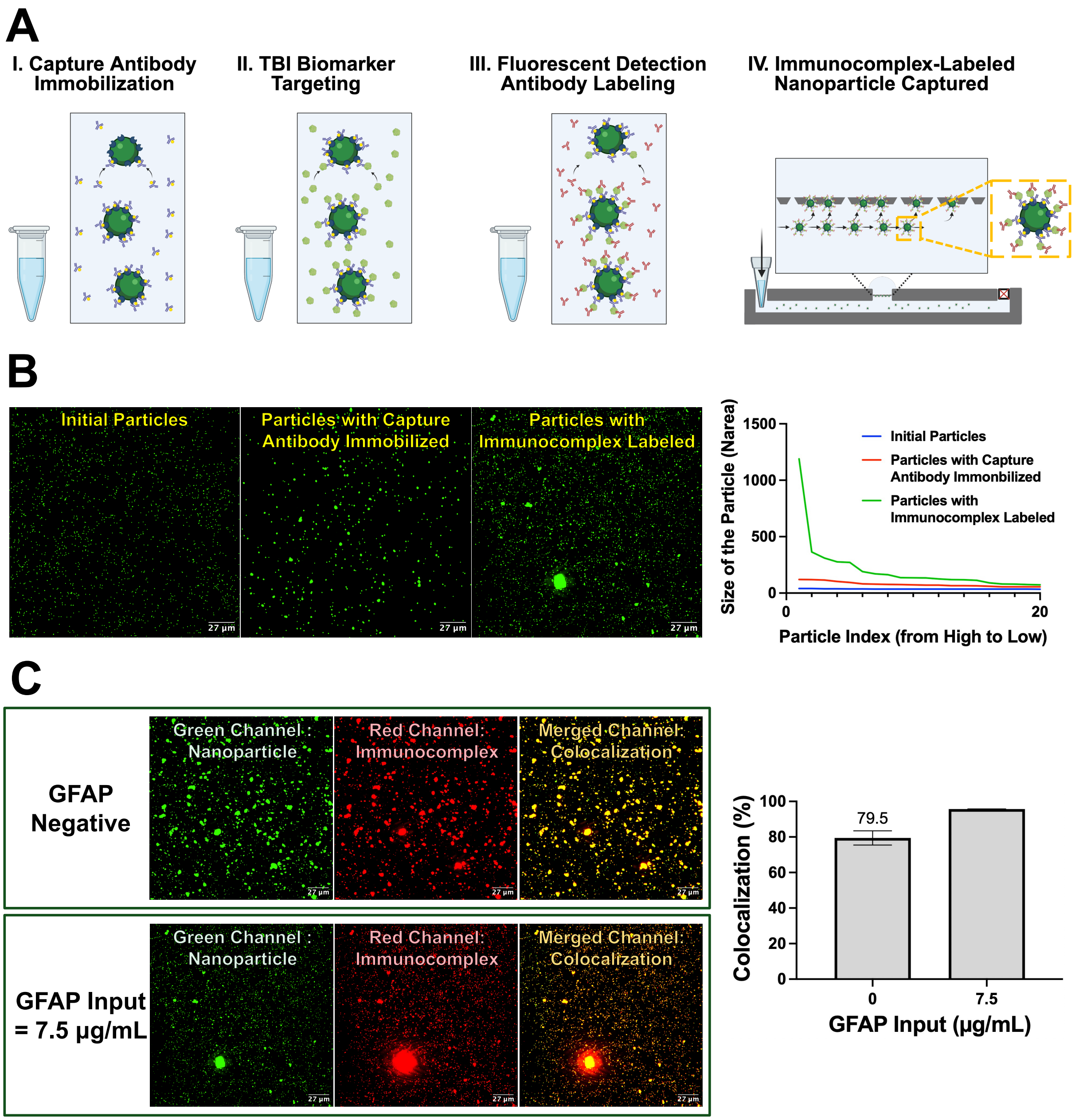


**Figure S1. Pre-Labeling of Nanoparticles Prior to Capture and Characterization. (A)** Pre-Labeling Workflow. All targeting and binding steps were performed in reaction tubes, followed by three 30 min washes with centrifugation after each step. The immunocomplex-labeled nanoparticles were then captured for signal detection. **(B)** Nanoparticle size characterization during sequential reagent labeling at GFAP = 7.5 µg/mL. Progressive aggregation of nanoparticles was observed, likely due to increased surface hydrophobicity from antibody and antigen binding. Particle size distributions were quantified using Narea, which accounts for both particle count and size without segmenting large particles. **(C)** Colocalization studies. High background fluorescence hindered signal discrimination. Fluorescence images (top row: GFAP = 0 µg/mL; bottom row: GFAP = 7.5 µg/mL) showed that the negative control exhibited a high colocalization rate (79.5%), likely due to nonspecific adsorption from altered surface properties of aggregated nanoparticles. The positive group displayed signal saturation, yielding a narrow detection window (~20%) between negative and positive signals (N = 3 fields of view per device).

**Movie M1 (in supplementary files).** Time-lapse confocal microscopy visualizing the controlled injection of functionalized nanoparticles and their real-time localization within the nanopores of the NPN membrane. The field of view shows a randomly selected region of the membrane, with nanoparticle injection initiated at the beginning of the recording. Individual nanoparticles appear within the field of view and rapidly become immobilized, remaining stationary without lateral movement or detachment, indicating successful capture by the nanomembrane and settling into the nanopores. **Movie M2 (in supplementary files).** Time-lapse confocal microscopy demonstrating the stability of nanoparticle hotspot capture under microfluidic flow. Nanoparticles in PBS ($2.5 \times 10^4$ nanoparticles/μL) was loaded into a syringe pump and injected through one open outlet at a flow rate of 60 μL/min, flushing the bottom channel of the device. The field of view shows that the captured nanoparticles remain firmly immobilized at the same locations during more than 1 minute of continuous flushing, confirming robust nanoparticle retention on the nanomembrane substrate.

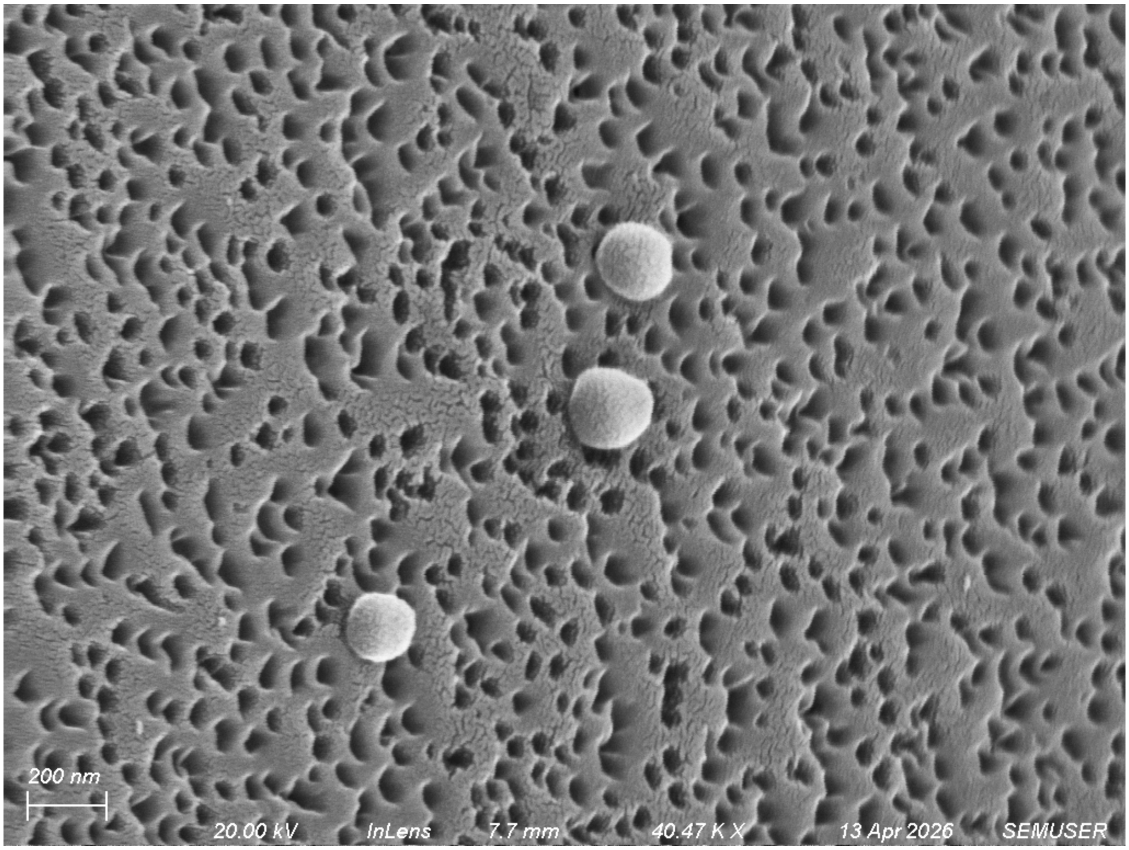


**Figure S2. Streptavidin-Labeled Fluorescent Nanoparticles Captured by the Nanomembrane as Target-Affnitiy Hotspots for Digital Immunoassay.** The SEM image showed 200 nm nanoparticles captured by the NPN (d ~ 60 nm) to form nanostructured surface with physically isolated and optically resolvable target-affinity ‘hotspots.’

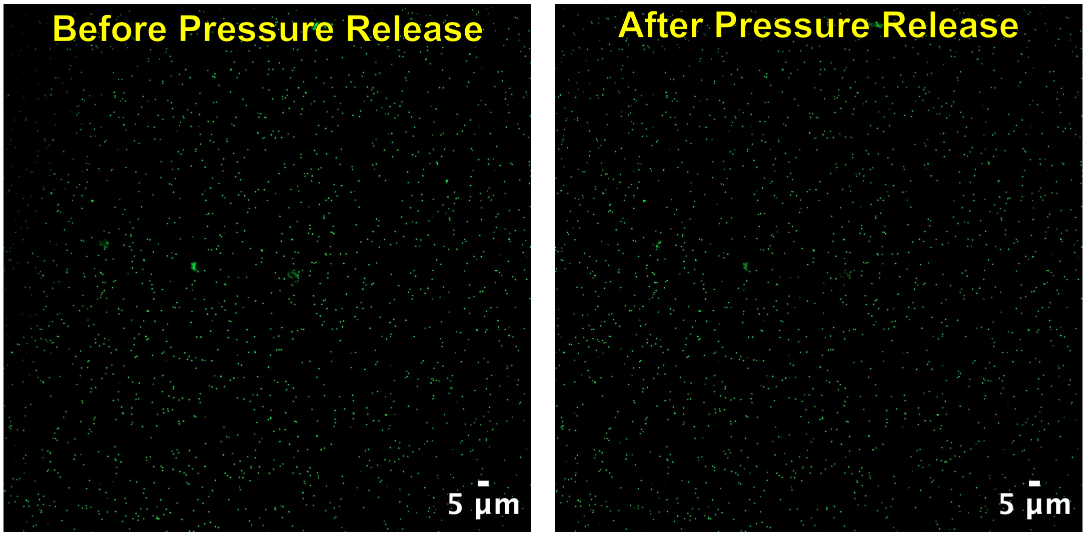


**Figure S3. Field of View at the end of Nanoparticle Injection. (Left)** Before pressure release, membrane deflection causes the field of view to appear out of focus. **(Right)** After pressure release, the membrane returns to its original position, restoring focus and enabling clear visualization of nanoparticles within the device.

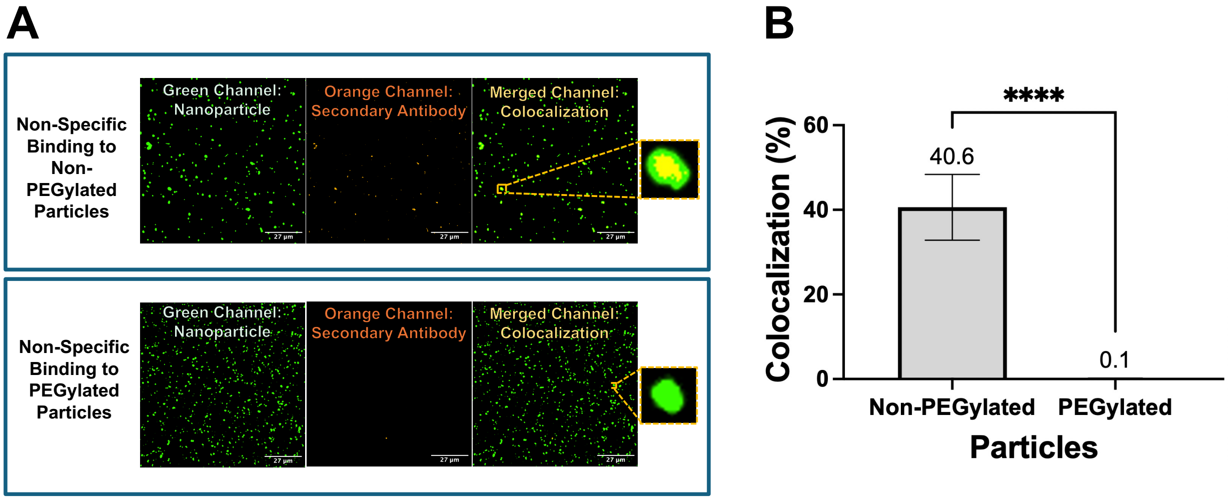


**Figure S4. Validation of PEGylated Nanoparticles as Specificity-Enhanced Substrates. (A)** Representative fields of view showing non-PEGylated and PEGylated nanoparticles after incubation with non-reactive Alexa Fluor 568–labeled anti-mouse secondary antibodies. PEGylated particles exhibited minimal colocalization, whereas non-PEGylated particles showed substantial nonspecific binding. **(B)** Quantification of nonspecific binding. PEGylated nanoparticles displayed only 0.1% colocalization with the secondary antibody compared to ~40% for non-PEGylated particles, demonstrating strong resistance to nonspecific interactions. Data are presented as mean ± SD (N = 6 fields of view per device). Statistical significance was determined by unpaired t-test (****P < 0.0001).

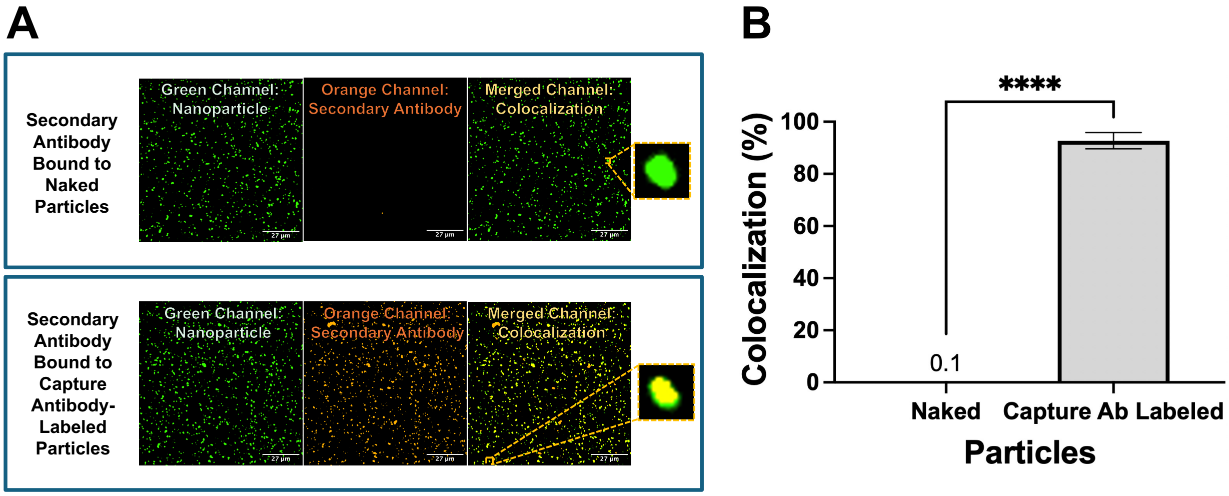


**Figure S5. Validation of PEGylated Nanoparticles as Effective Reagent Targeting Substrate. (A)** Representative fields of view of initial nanoparticles and capture antibody–conjugated PEGylated nanoparticles after incubation with the secondary antibody. Capture antibody-labeled particles showed significant colocalization, indicating preserved antibody accessibility and conjugation efficiency after PEG spacer incorporation. **(B)** Quantitative analysis of capture antibody conjugation efficiency. Incubation of Alexa Fluor 568–labeled anti-mouse secondary antibody with capture antibody-labeled PEGylated nanoparticles resulted in 92.7% colocalization, confirming successful capture antibody functionalization. Data are presented as mean ± SD (N = 6 fields of view per device). Statistical significance was determined by unpaired t-test (****$P < 0.0001$).

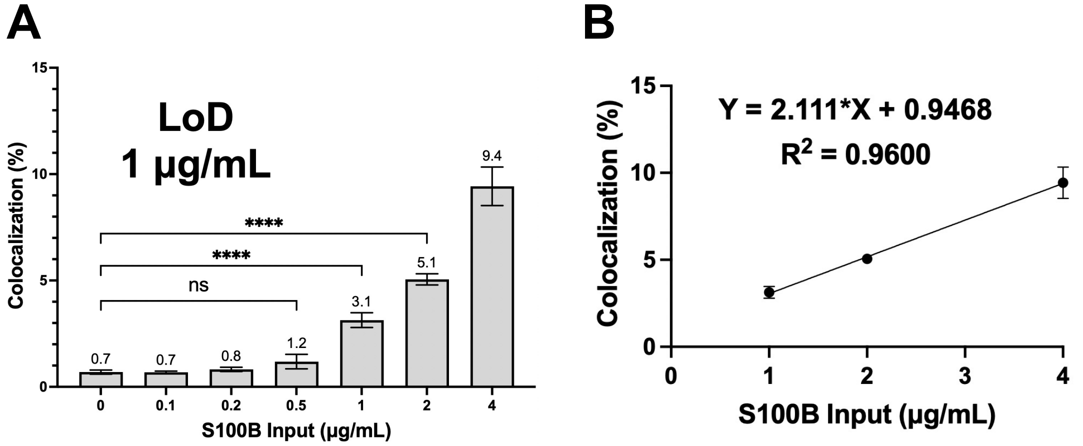


**Figure S6. CAD-IA for S100B Detection in 10% FBS.** The assay showed consistent responses with a LoD of 1 µg/mL and a linear range from 1 to 4 µg/mL ($R^2$ = 0.9600). Data are presented as mean ± SD (N = 6 fields of view per device). Statistical significance was assessed using ordinary one-way ANOVA followed by Dunnett's multiple comparisons test. ****p < 0.0001. Statistical significance is primarily reported for concentrations near the LoD, as higher concentrations exhibited clear separation from the background. Large error bars near the blank reflect Poisson counting statistics in the low-count regime, where small numbers produce high relative variance.

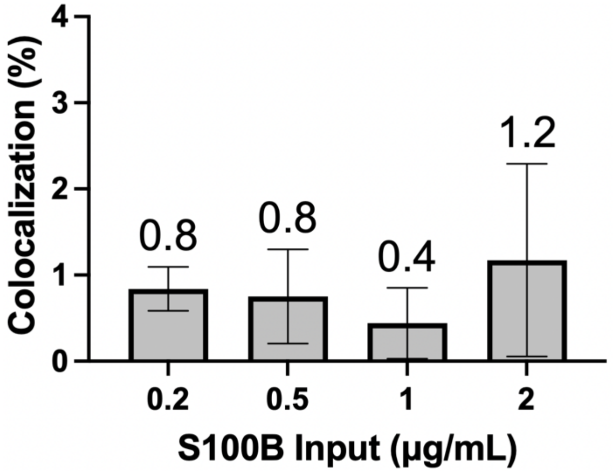


**Figure S7. CAD-IA for S100B Detection in 100% FBS.** CAD-IA in 100% FBS showed signal suppression to baseline across all antigen inputs, likely due to nanoparticle surface fouling by matrix components, which hindered target accessibility. Data are presented as mean ± SD (N = 6 fields of view per device). Large error bars near the blank reflect Poisson counting statistics in the low-count regime, where small numbers produce high relative variance.

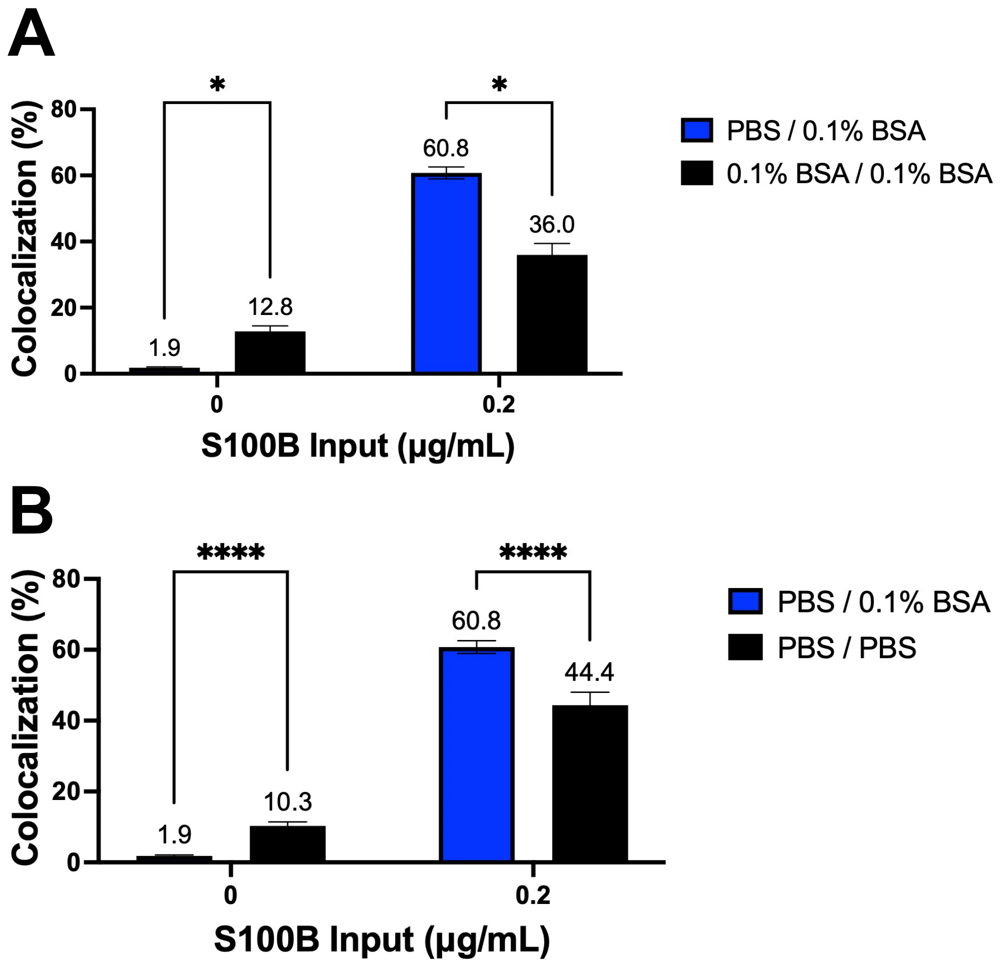


**Figure S8. Optimization of Solvent System for TSA-Modified CAD-IA. (A)** Evaluation of solvent systems for the capture antibody. Dissolving the capture antibody in PBS produced significantly stronger signals compared to 0.1% BSA, indicating improved surface immobilization efficiency. Statistical significance was assessed using multiple unpaired *t* tests (*$p < 0.000001$). **(B)** Evaluation of solvent systems for the detection antibody. In contrast, dissolving the detection antibody in 0.1% BSA significantly reduced background signal, confirming the role of BSA as a blocking reagent that improves the signal-to-noise ratio. Data are presented as mean ± SD (N = 6 fields of view per device). Statistical significance was assessed using multiple unpaired *t* tests (****$p < 0.0001$). Legend labels denote solvent combinations for capture/detection antibodies; for example, "PBS / 0.1% BSA" indicates that the capture antibody was dissolved in PBS and the detection antibody in 0.1% BSA / PBS.

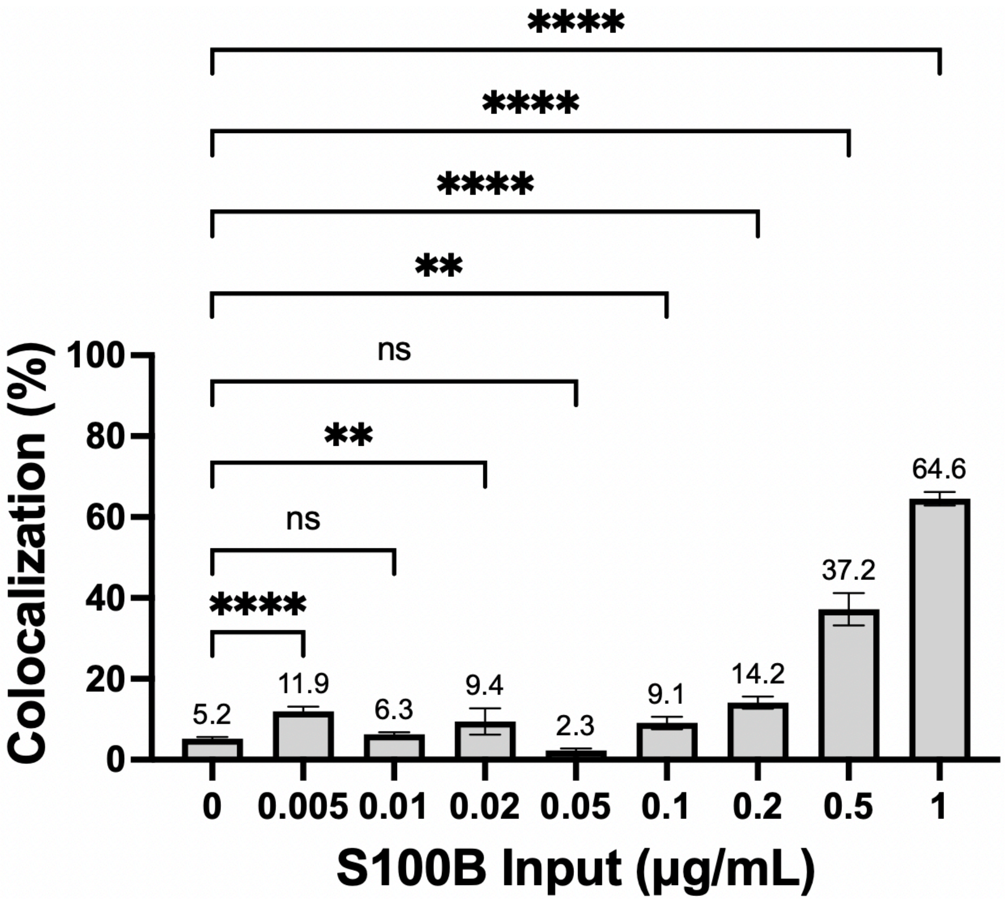


**Figure S9. Effect of Ultracentrifugation on Reagent Performance.** Non-ultracentrifuged reagents produced strong and consistent signals, whereas ultracentrifuged antibodies exhibited weakened and more variable responses, indicating potential antibody loss and reduced functional activity following ultracentrifugation. Data are presented as mean ± SD (N = 6 fields of view per device). Statistical significance was evaluated using ordinary one-way ANOVA with Dunnett's multiple comparisons test (****p < 0.0001; **p = 0.0041 for 0 vs 0.02 µg/mL; **p = 0.0092 for 0 vs 0.1 µg/mL).

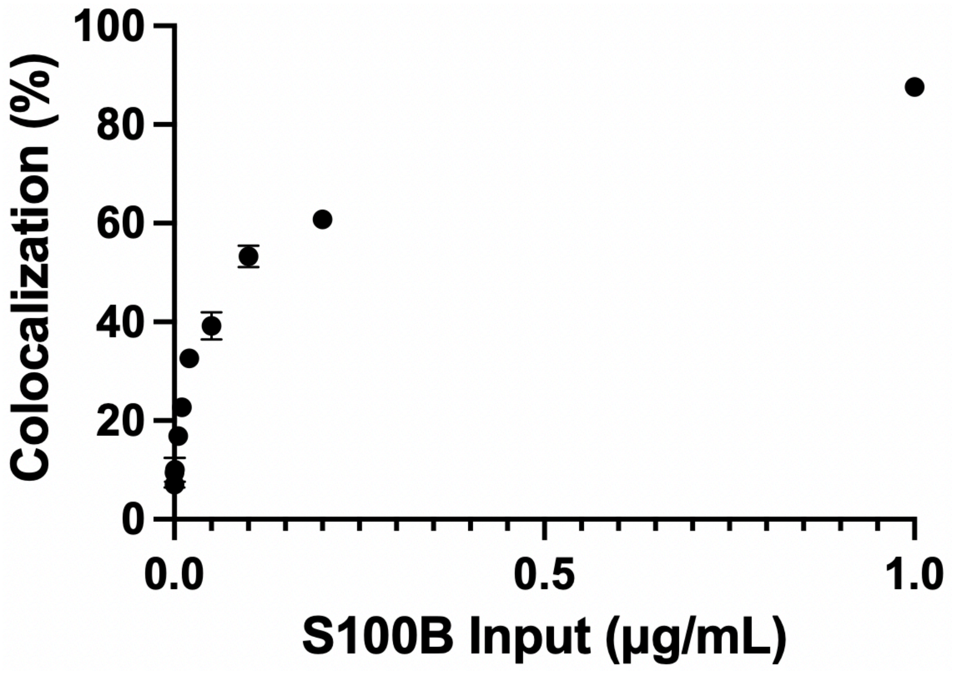


**Figure S10. Nonlinear Dose–Response Behavior Induced by Enzymatic Amplification.** In contrast to the linear dose–response observed in the non-amplified assay, enzymatic signal amplification produces a nonlinear relationship between fluorescence signal and analyte concentration. Data are presented as mean ± SD (N = 6 fields of view per device).
